# Revisiting the role of the spindle assembly checkpoint in the formation of gross chromosomal rearrangements in *Saccharomyces cerevisiae*

**DOI:** 10.1101/2024.04.11.589040

**Authors:** Yue Yao, Ziqing Yin, Fernando R. Rosas Bringas, Jonathan Boudeman, Daniele Novarina, Michael Chang

**Author notes:** These authors contributed equally (Yue Yao, Ziqing Yin).

## Abstract

Multiple pathways are known to suppress the formation of gross chromosomal rearrangements (GCRs), which can cause human diseases including cancer. In contrast, much less is known about pathways that promote their formation. The spindle assembly checkpoint (SAC), which ensures the proper separation of chromosomes during mitosis, has been reported to promote GCR, possibly by delaying mitosis to allow GCR-inducing DNA repair to occur. Here we show that this conclusion is the result of an experimental artifact arising from the synthetic lethality caused by disruption of the SAC and loss of the *CIN8* gene, which is often lost in the genetic assay used to select for GCRs. After correcting for this artifact, we find no role of the SAC in promoting GCR.

**Significance statement:** A gross chromosomal rearrangement (GCR) is an abnormal structural change of a native chromosome. Examples of GCRs include deletions, duplications, inversions, and translocations. GCRs can lead to genetic diseases such as cancer. A previous study implicated the spindle assembly checkpoint (SAC), which ensures the proper separation of chromosomes during cell division, in facilitating the formation of GCRs. In this study, we show that this is not the case; the SAC does not promote GCR.

## Introduction

Gross chromosomal rearrangements (GCRs) are large-scale changes in the structure of chromosomes. GCRs, which include interstitial deletions, duplications, inversions, and translocations, can affect the number, position, and orientation of genes within a chromosome or between chromosomes. They can occur spontaneously during cell division or as a result of exposure to environmental factors such as radiation or chemical mutagens. GCRs are associated with several genetic diseases, are frequently observed in cancer cells, and can contribute to the initiation or progression of cancer (1, 2) .

The mechanisms that suppress the formation of GCRs have been best studied in the budding yeast *Saccharomyces cerevisiae* using genetic assays, such as the “classical” GCR assay developed by Chen and Kolodner, and variations of this assay (3, 4). In the classical GCR assay, two counterselectable markers, *URA3* and *CAN1*, are located on the left arm of chromosome V between the telomere and *PCM1*, the most telomere-proximal essential gene. A GCR involving the loss of both markers renders the cell resistant to 5-fluoroorotic acid (5-FOA) and canavanine. Using these assays, many GCR suppressing pathways have been identified. These pathways are involved in processes such as DNA replication and repair, S-phase checkpoints, chromatin assembly, telomere maintenance, oxidative stress response, and suppression of R-loop accumulation (4).

Several pathways are also known to promote GCR formation. Among these, de novo telomere addition, nonhomologous end-joining (NHEJ), and homologous recombination (HR) are notably well-characterized (4). De novo telomere addition occurs when a broken chromosome end is healed by the addition of a new telomere, resulting in truncation of the chromosome. NHEJ and HR are the two main pathways for the repair of double-strand breaks, but inappropriate NHEJ and HR can lead to translocations or interstitial deletions. However, deletion of genes important for NHEJ and HR often do not reduce, and can even increase, the rate of GCRs, because NHEJ and HR act to both suppress and generate GCRs (4). In addition, transcription can promote GCR, likely due to transcription-dependent replication stress (5). The Rad1-Rad10 endonuclease also promotes GCR, but how it does so remains unclear, with multiple mechanisms proposed (5, 6). Lastly, the spindle assembly checkpoint (SAC), the Bub2-Bfa1 GTPase activating protein complex, and the Ctf18-Dcc1-Ctf8-RFC complex have all been implicated in GCR formation induced by various genetic mutations (7). The SAC ensures accurate chromosome separation during mitosis by delaying the metaphase/anaphase transition until all kinetochores are attached to microtubules (8); the Bub2-Bfa1 complex prevents premature mitotic exit (9); and the Ctf18-Dcc1-Ctf8-RFC complex is important for preventing chromosome loss and precocious sister chromatid separation (10). It was proposed that DNA lesions that lead to GCR activate the SAC and delay mitotic exit, allowing time for GCR-inducing repair to occur; without this cell cycle delay, cells would progress through mitosis before the damage can be repaired, causing increased lethality and an apparent suppression of GCRs (7).

To explore the impact of interstitial telomeric sequences (ITSs) on GCR, we modified this assay by inserting a 50-bp ITS between *PCM1* and *CAN1* (Figure 1A). This modification results in a >1000-fold increase in the GCR rate (Rosas Bringas and Yin et al., accompanying manuscript). Subsequently, we performed a genome-wide screen and identified genes that promote ITS-induced GCR, including SAC genes, *BUB2* and *BFA1*, and *CTF8* and *DCC1*, consistent with the previous finding that these genes play a role in GCR formation (7). However, we find that the apparent GCR-suppressing effect of these mutants can be attributed to the known synthetic lethality arising from the deletion of any of these genes combined with the loss of *CIN8* (11, 12), which encodes a bipolar kinesin motor protein that plays a pivotal role in mitotic spindle assembly and chromosome segregation (13, 14). *CIN8* is located immediately downstream of the inserted ITS (Figure 1A), and is often lost during GCR formation in the classical GCR assay. We find that SAC, *bub2Δ, bfa1Δ, ctf8Δ*, and *dcc1Δ* mutants do not suppress GCRs in strains with an additional copy of *CIN8* located elsewhere in the genome. Therefore, we conclude that the SAC, Bub2-Bfa1, and the Ctf18-Dcc1-Ctf8-RFC complex do not significantly contribute to GCR formation.

**Figure 1.**
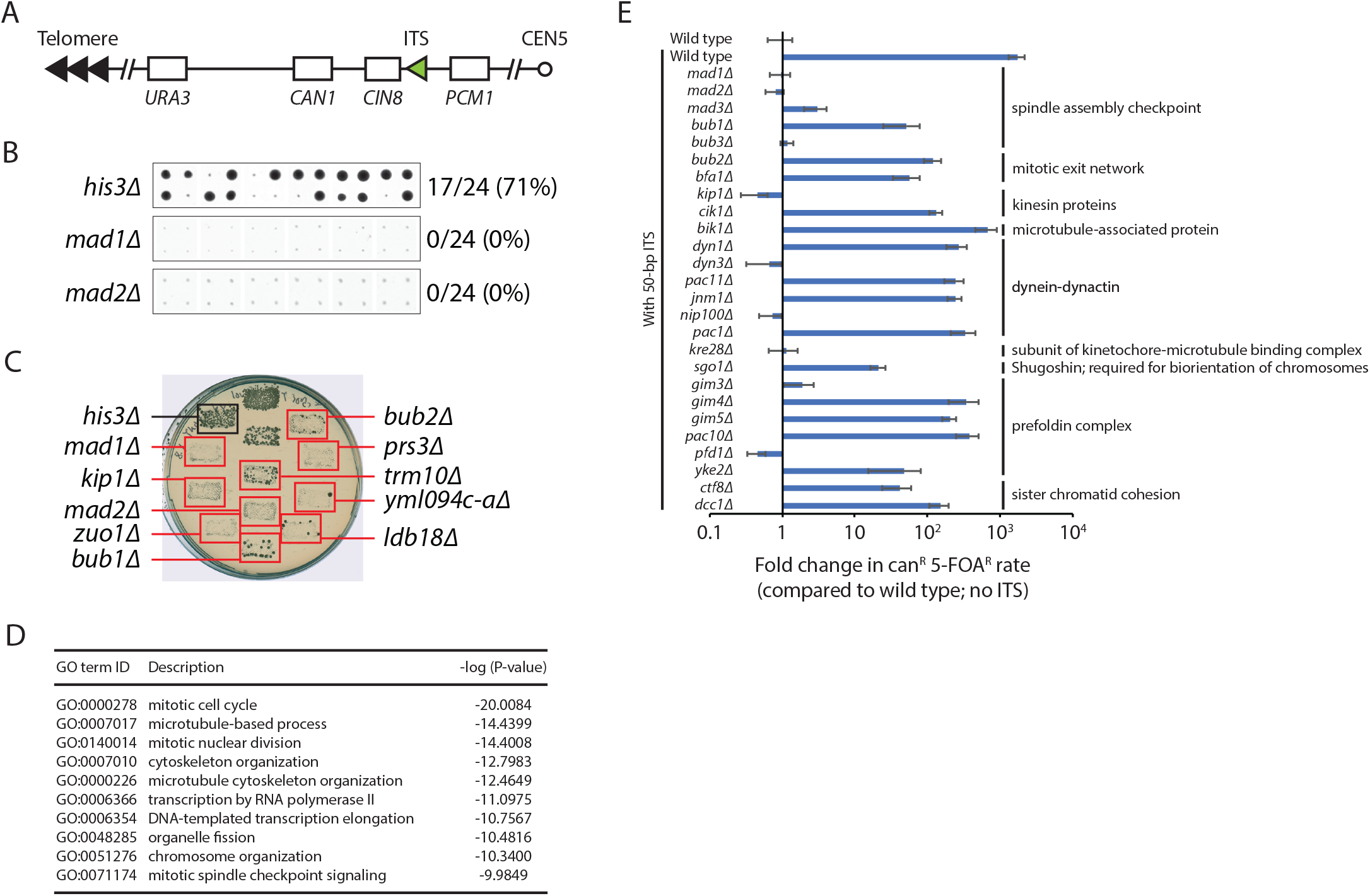
A genome-wide screen for genes that promote the formation of ITS-induced GCRs identifies genes with functions in microtubule-based processes and chromosome segregation. (**A**) Schematic diagram of the ITS-GCR assay. A GCR that leads to the simultaneous loss of two genetic markers, *URA3* and *CAN1*, can be selected by growth on 5-FOA and canavanine. A 50-bp ITS was inserted between *CIN8* and *PCM1*, the most telomere-proximal essential gene on the left arm of chromosome V. (**B**) A high-throughput screen was performed (Rosas Bringas and Yin et al., accompanying manuscript). All 24 replica-pinned colonies on media containing both canavanine and 5-FOA of the *his3Δ* control strain and two selected mutants with decreased GCR frequencies are shown. (**C**) Putative hits were tested in a patch-and-replica-plate assay. An example plate is shown. Hits that tested positive are indicated by red boxes. A negative control (*his3Δ*) and a positive control (*bub2Δ*) were included on each plate. (**D**) The top 10 GO terms enriched in the hits that tested positive in the patch-and-replica-plate assay are shown. (**E**) Fold change in canavanine/5-FOA-resistance rate of the indicated strains, relative to the wild-type strain without an ITS, is plotted. Error bars represent SEM (n = 3–6).

## Results and Discussion

### A genome-wide screen for genes that promote the formation of ITS-induced GCRs identifies spindle assembly checkpoint and mitotic exit genes

To identify genes that promote the formation of ITS-induced GCRs, we modified the classical GCR assay (3) by inserting a 50 bp-ITS between the most telomeric essential gene on the left arm of chromosome V (*PCM1*) and two counterselectable markers (*CAN1* and *URA3*) (Figure 1A). We used this ITS-GCR reporter to screen the yeast knockout (YKO) and conditional temperature-sensitive (ts) strain libraries (Rosas Bringas and Yin et al., accompanying manuscript). A total of 213 YKO and 93 ts hits were identified in the screen (Figure 1B) and confirmed in a patch-and-replica-plate assay (Figure 1C). The hits are enriched for genes involved in the mitotic cell cycle and microtubule-based processes, including the SAC (Figure 1D).

To further validate the hits, we performed fluctuation tests for a subset of mutants. We find that deletion of genes important for the SAC (*MAD1, MAD2, MAD3, BUB1, BUB3*), the Bub2-Bfa1 complex, and the Ctf18-Dcc1-Ctf8-RFC complex (*CTF8, DCC1*) reduces the increased GCR rate caused by the ITS (Figure 1E), reminiscent of the previous finding that deletion of these genes can suppress GCRs in mutants with elevated GCR rates, as assayed using the classical GCR assay (7). Additionally, we find that deletions of many other genes with microtubule-related functions—including those that encode kinesin and microtubule-associated proteins (*KIP1, CIK1, BIK1*), dynein-dynactin (*DYN1, DYN3, PAC11, JNM1, NIP100, PAC1*), proteins involved in the attachment of microtubules to kinetochores (*KRE28, SGO1*), and subunits of the prefoldin complex (*GIM3, GIM4, GIM5, PAC10, PFD1, YKE2*), which is important for microtubule biogenesis—also decrease the ITS-induced GCR rate (Figure 1E).

### Defects in the SAC or Bub2-Bfa1 cannot suppress ITS-induced GCR rate when an extra copy of *CIN8* is present

While investigating the mechanism by which these genes could promote GCR, we realized that all the gene deletions shown in Figure 1E have been reported to be synthetic lethal with co-deletion of the *CIN8* gene (11, 12, 15, 16), which encodes a kinesin motor protein (13, 14) that is often lost along with *CAN1* and *URA3* when selecting for GCRs using the classical GCR assay (Figure 1A). Thus, the apparent suppression of GCRs by these mutants could be explained by the inability of these mutants to survive a GCR event that also results in the loss of *CIN8*. To examine the real effect of the SAC and the Bub2-Bfa1 complex on GCR formation, we determined the GCR rate of diploid strains that have one chromosome V containing the ITS-GCR reporter (*URA3, CAN1*, and the ITS) while the homologous chromosome V does not (Figure 2A). Importantly, in this setting, any synthetic lethality caused by loss of *CIN8* is circumvented by the presence of another copy of *CIN8* on the homologous chromosome. We first tested a wild-type diploid strain containing the 50-bp ITS and observed that the GCR rate does not significantly change compared to the wild-type haploid containing the 50-bp ITS. We then tested heterozygous and homozygous SAC and *bub2*/*bfa1* mutants; we find that disruption of the SAC or the Bub2-Bfa1 complex does not cause a reduction in GCR rate. In fact, *bub1Δ/bub1Δ* and *bub3Δ/bub3Δ* diploid strains exhibit an apparent 77-fold and 117-fold increase in GCR rate, respectively, compared to the wild-type diploid (Figure 2A).

**Figure 2.**
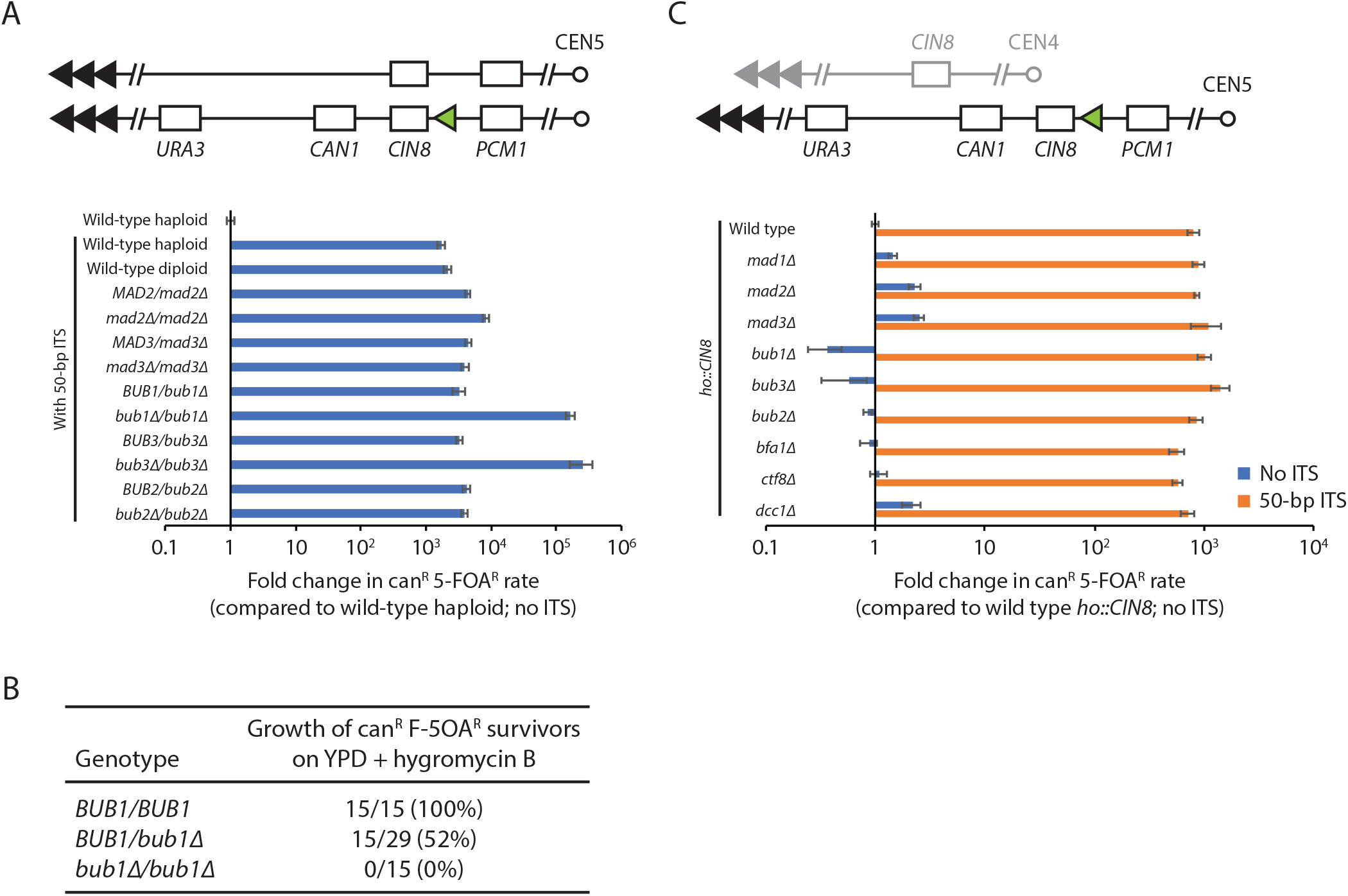
Defects in the SAC, the Bub2-Bfa1 complex, or sister chromatid cohesion cannot suppress ITS-induced GCR rate when an extra copy of *CIN8* is present. (**A**) Fold change in canavanine/5-FOA-resistance rate of the indicated diploid strains, relative to the wild-type haploid strain without an ITS, is plotted. The diploid strains have one chromosome V with the ITS-GCR reporter, while the homologous chromosome V does not. (**B**) The percentages of canavanine/5-FOA-resistant survivors, generated from the *BUB1/BUB1, BUB1Δ/bub1Δ*, and *bub1Δ/bub1Δ* strains in **A**, that are also resistant to hygromycin B are shown. (**C**) Fold change in GCR rate of the indicated strains, which all have an extra copy of *CIN8* inserted at the *ho* locus, is plotted. Fold changes are relative to the wild-type haploid strain with the extra copy of *CIN8* gene, but without an ITS. Error bars represent SEM (n = 3–6).

Bub1 (likely together with its partner Bub3) has a SAC-independent function to recruit Sgo1 to kinetochores, which is important for ensuring that sister kinetochores are attached to microtubules from opposite poles (17). Cells lacking Bub1 or Bub3 missegregate chromosomes at a higher rate than other SAC mutants due to the persistence of uncorrected syntelic attachments (18, 19). Thus, the apparent increase in GCR rate in *bub1Δ/bub1Δ* and *bub3Δ/bub3Δ* strains could be due to loss of the *URA3*-and *CAN1*-containing chromosome. Such an increase would not be apparent in a haploid setting because loss of the sole copy of chromosome V would be lethal, whereas monosomy in an otherwise diploid yeast strain is not (20). Since an *hphMX* cassette, which provides resistance to hygromycin B, is integrated on the centromeric side of the ITS, and because practically all GCRs selected in this assay involve a de novo telomere addition at the ITS that leaves the *hphMX* cassette in place (Rosas Bringas and Yin, accompanying manuscript), we can use hygromycin B resistance to differentiate between a survivor of a GCR event (hygromycin B resistant) from a survivor of a chromosome loss event (hygromycin B sensitive). We tested canavanine-and 5-FOA-resistant survivors derived from *BUB1/BUB1, BUB1/bub1Δ*, and *bub1Δ/bub1Δ* strains and found that all *BUB1/BUB1* survivors were resistant to hygromycin B while none of the *bub1Δ/bub1Δ* survivors were, indicating that the increase in canavanine-and 5-FOA-resistant *bub1Δ/bub1Δ* survivors is a result of an increase in chromosome loss, not GCR (Figure 2B). Interestingly, although there is no increase in the rate of canavanine-and 5-FOA-resistance for the *BUB1/bub1Δ* strain (Figure 2A), approximately half of the *BUB1/bub1Δ* survivors were resistant to hygromycin B (Figure 2B), indicating Bub1 haploinsufficiency increases loss of chromosome V to levels similar to the GCR rate obtained using the 50-bp ITS-GCR assay.

We further assessed the role of the SAC, the Bub2-Bfa1 complex, and the Ctf18-Dcc1-Ctf8 complex in the formation of GCRs by using haploid strains, with and without the 50-bp ITS, that have an extra copy of *CIN8* integrated at the *ho* locus on chromosome IV. In this setting, deletion of genes involved in these processes/complexes causes only mild (less than threefold) changes in GCR rate (Figure 2C). Taken together, our results indicate that the SAC, along with the Bub2-Bfa1 and Ctf18-Dcc1-Ctf8 complexes, neither promote nor suppress the formation of GCRs.

Our findings highlight an important point to consider when using genetic assays. If the assay results in the loss of genes not directly related to the assay, it is important to consider whether a synthetic lethal genetic interaction exists between one of these genes and any mutant being studied using the assay. If such a synthetic lethal interaction exists, it (1) may give the false appearance that the mutant decreases the rate of the studied genetic event, and (2) may mask an actual increase in the rate. For the classical GCR assay, there are 23 open reading frames (several of which are classified as dubious in the *Saccharomyces* Genome Database) between *PCM1* and the telomere on the left arm of chromosome V. While loss of any of these genes may pose such a problem, most of the known genetic interactions for this group of genes involves *CIN8*. Other assays involving other regions of the genome will be affected by a different set of synthetic lethal interactions.

## Materials and methods

### Yeast strains and plasmids

All yeast strains used in this study are listed in Table S1. Standard yeast genetic and molecular methods were used (21, 22). Strains containing an extra copy of *CIN8* were constructed as follows: *CIN8* with its own promoter and terminator was PCR-amplified from yeast genomic DNA and cloned via BsaI Golden Gate assembly into plasmid pYTK164 (GFP-dropout *HO*-locus integration vector constructed with the MoClo-YTK (23). The resulting plasmid (pDN60.3) was digested with NotI for integration of *CIN8* at the *ho* locus.

### High-throughput replica pinning screen

The high-throughput replica-pinning screen was performed essentially as previously described (24). Details can be found in the accompanying manuscript (Rosas Bringas and Yin et al.).

### Fluctuation tests of GCR rates

Fluctuation tests for the quantification of GCR rates were performed essentially as previously described (25) by transferring entire single colonies from YPD plates to 4 ml of YPD liquid medium and grown to saturation. 50 µl of a 10^5^-fold dilution was plated in YPD plates. An strain-dependent quantity of cells was plated on SD-arg+canavanine+5-FOA. Colonies were counted after incubation at 30ºC for 3-5 days. The number of GCR (can^R^5-FOA^R^) colonies was used to calculate the GCR rate by the method of the median (26).

### Gene ontology enrichment analysis

The GO term finder tool (http://go.princeton.edu/) was used to query biological process enrichment for each gene set, with a P-value cutoff of 0.01 and Bonferroni correction applied. REVIGO (27) was used to further analyze the GO term enrichment data, using the “Medium (0.7)” term similarity filter and the simRel score as semantic similarity measure. As a result, terms with a frequency more than 10% in the REVIGO output were eliminated for being too broad.

## Acknowledgments and funding sources

We thank A. Milias-Argeitis for providing plasmid pYTK164. Y.Y. was supported by an Abel Tasman Talent Program scholarship from the University of Groningen. Z. Yin was supported by a scholarship from the Nanjing Huimou Medi-Tech Co. F.R.R.B. was supported by a Consejo Nacional de Ciencia y Tecnología (CONACYT) scholarship. Work in the laboratory of M. Chang was supported by an Open Competition M-2 grant from the Dutch Research Council.

---

**Table S1.**
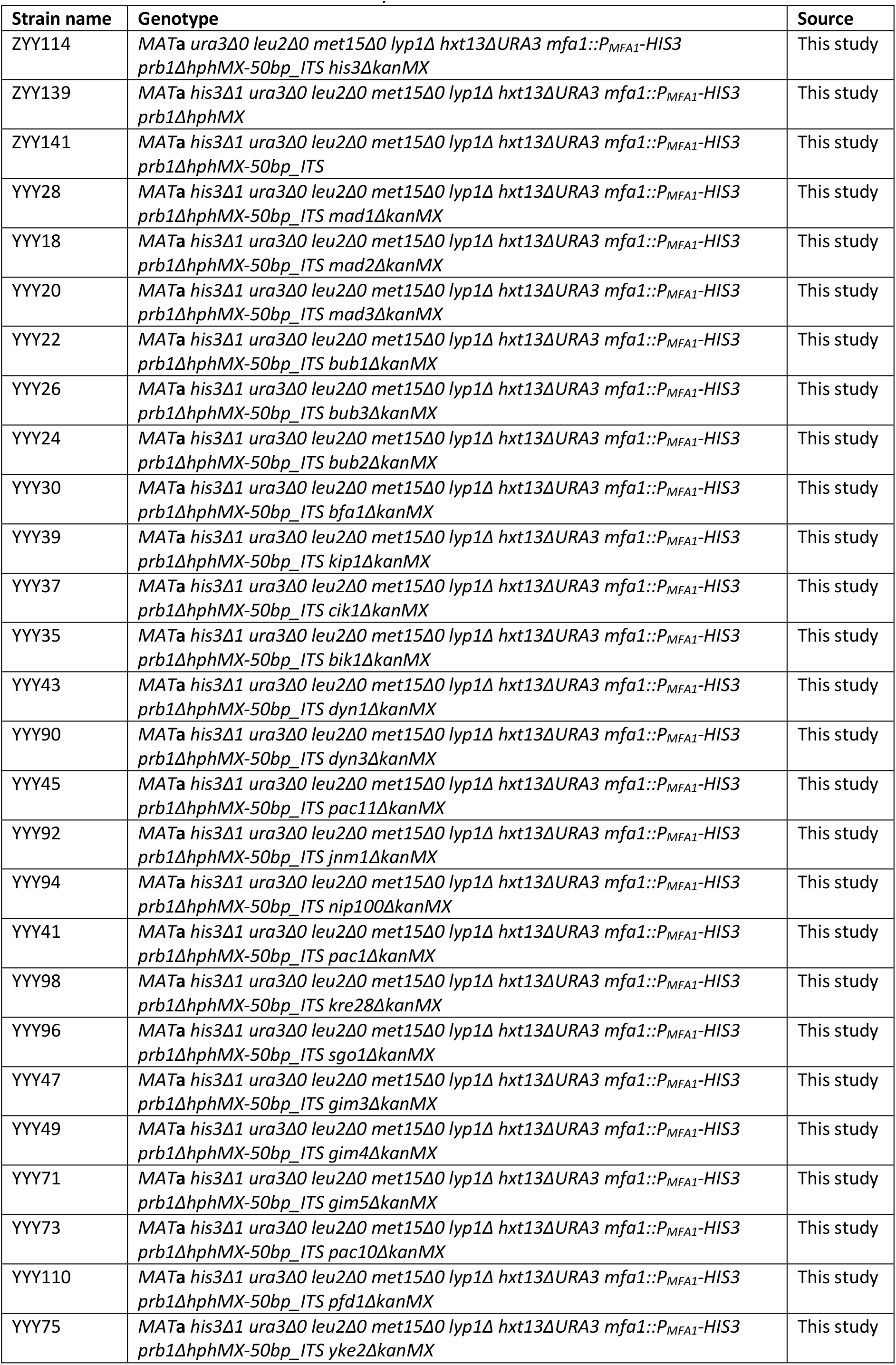
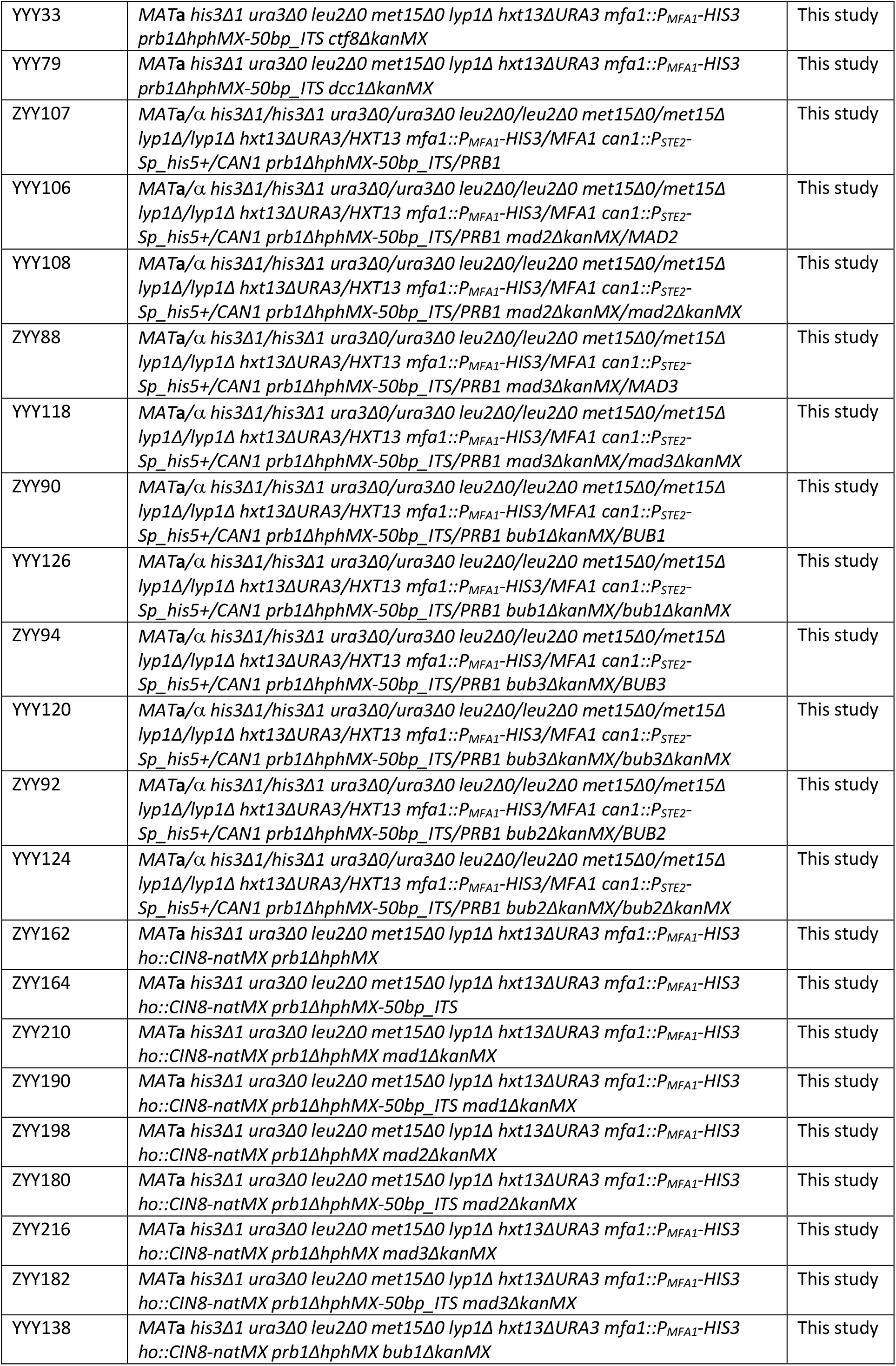
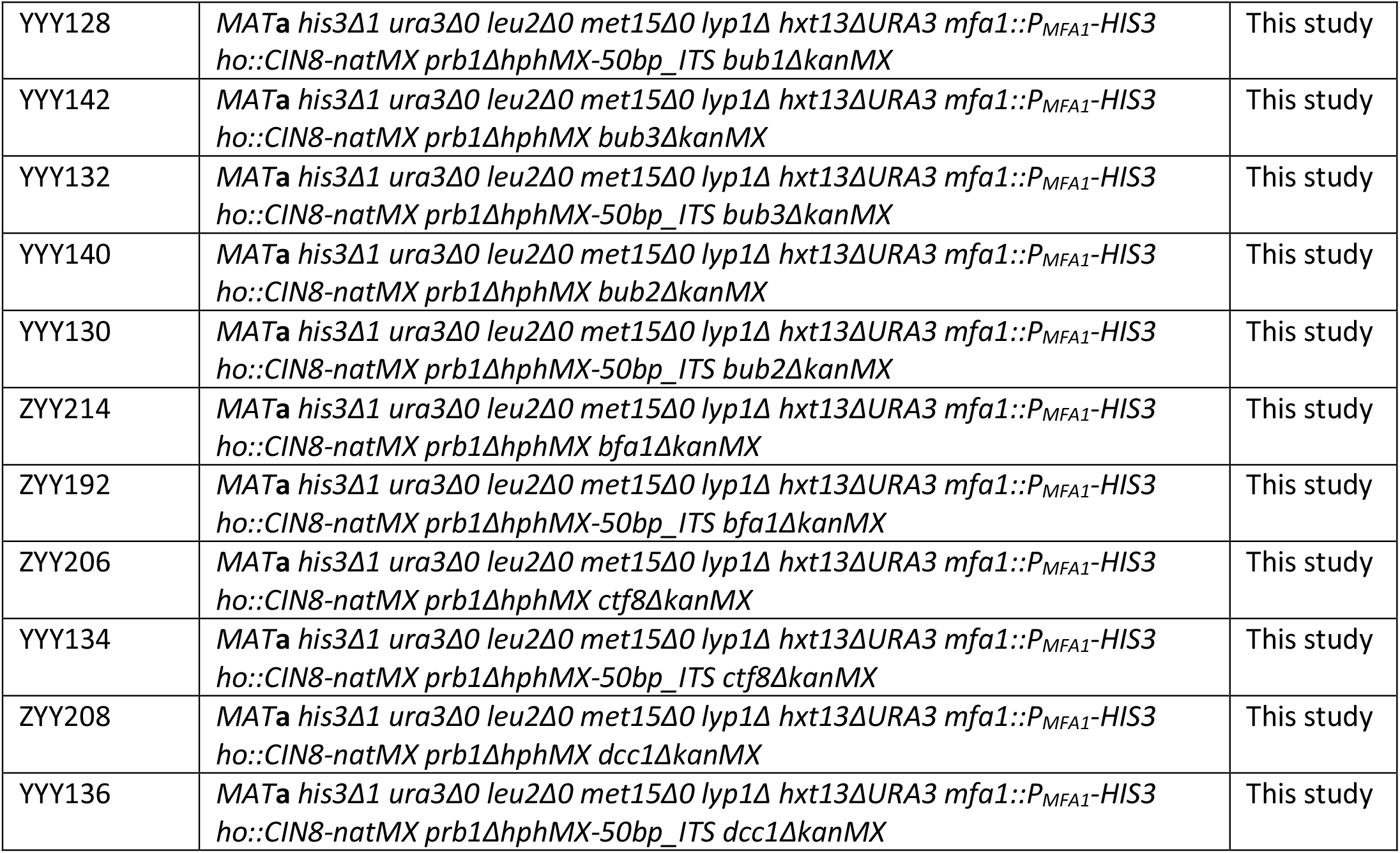
Yeast strains used in this study.

